# *Rattus economicus*: A neural model for rational regulation of water-seeking effort in rodents

**DOI:** 10.1101/2021.03.01.433467

**Authors:** Pamela Reinagel

## Abstract

In the laboratory, animals’ motivation to work tends to be positively correlated with reward magnitude. But in nature, rewards earned by work are essential to survival (e.g., working to find water), and the payoff of that work can vary on long timescales (e.g., seasonally). Under these constraints, the strategy of working less when rewards are small could be fatal. We found that instead, rats in a closed economy did more work for water rewards when the rewards were stably smaller, a phenomenon also observed in human labor supply curves. Like human consumers, rats also showed elasticity of demand, consuming far more water per day when its price in effort was lower. The neural mechanisms underlying such “rational” market behaviors remain largely unexplored. We propose a dynamic utility maximization model that can account for the dependence of rat labor supply (trials/day) on the wage rate (ml/trial), and also predict the temporal dynamics of when rats work. Based on data from mice, we hypothesize that SFO^GLUT^ neurons in *lamina terminalis* continuously compute the instantaneous marginal utility of voluntary work for water reward, and causally determine the amount and timing of work.

## Introduction

When animals have two ways to get a resource like water, they tend to choose the way that gets them more water for less work. Neural mechanisms underlying choices involving value comparisons are well-studied (*1, 2*). The reward literature has focused on how the relative subjective value or “utility” of each option is determined by weighing benefits (such as reward magnitude or quality) against costs (such as delay, risk, or effort). The identified neural mechanisms for utility computation mostly involve striatal and limbic reward circuits and dopamine.

Much less is known about how animals assess the absolute value of a single available option to decide whether or not to attempt to harvest a potential reward. In one of the few such studies, when mice were offered only one way to get water at a time, they worked harder during the time blocks when the water reward was larger (*3*). This makes sense – save energy for when the work will pay off most – but it can’t be the whole story. If motivation were driven entirely by expected reward, animals would be less motivated to work for water during a drought (because they would expect less reward per unit of effort) and might die of thirst. This problem is partly offset by the fact that the perceived value of a reward is normalized according to recent experience, such that rewards that would have been considered small in a rich environment are perceived as large relative to a lean environment (*4-6*). But normalization would at best equalize motivation between rich and lean environments. If the difficulty of getting water changes slowly compared to the timescale of physiological necessity, animals must invest the most effort to gain it precisely when the reward for that effort is least.

To explore how animals adapt to this kind of challenge, we maintained rats in a live-in environment in which all their water was earned by performing a difficult sensory task. We varied the reward magnitude and measured rats’ effort output and water consumption. As expected, rats did more trials per day when the reward per trial was smaller, thus maintaining healthy hydration levels regardless of reward size. More surprisingly, however, rats worked for more water per day (and far more than they needed) when it was easier to earn. This suggests that they can regulate their consumption dramatically (up to three-fold) to conserve effort when times are lean or cash in during times of abundance. In economic terms, rats show strong elasticity of demand for water, even though essential commodities without substitutes are expected to be inelastic.

Classic animal behavior studies noted both these effects in experiments designed to validate economic utility maximization theory (*7-9*). Here we revisit and extend that theoretical framework with the goal of relating utility maximization to behavioral dynamics and candidate neural mechanisms. This study differs from the recent literature on utility maximization in choice behavior in two ways. Behaviorally, we focus here on the choice between action and inaction under a closed economy with closed-loop feedback on value (where state changes as a function of past choices). Mechanistically, we implicate *lamina terminalis*, a forebrain circuit which has not been previously linked to utility computations.

## Results

### Experimental approach

Rats performed a visual discrimination task to earn water rewards (*10, 11*). Briefly, an operant chamber was connected to each rat’s cage as the rat’s sole source of water. Rats could enter the chamber and initiate a trial at any time, upon which a visual motion stimulus was displayed on one wall of the chamber. The direction of motion indicated which of the two response ports would deliver a water reward; a response at the other port resulted in a brief time-out. The visual discrimination was difficult enough that rats made errors. We know rats consider this “work” because they will not do trials if there is no water reward, or if they are not thirsty. The reward volume was held constant for blocks of 1-2 weeks, and varied between time blocks. Below we use the term “reward size” to refer to the *a priori* expected reward of a trial (the volume of one reward multiplied by the probability of earning a reward). This value was stable for days and therefore known to the rat at the time of each decision to initiate a trial. We measured how many trials a rat performed and how much water reward a rat consumed per day. For further experimental details see Methods.

### Observations motivating the model

We measured the steady-state trial rate of N=4 rats which had access to the task 24 hours/day in their home cage and were tested for at least eight non-overlapping steady-state time blocks with reward sizes spanning at least a 50µl range, allowing us to ask how behavior correlated with reward size. Trial rate *L* declined with increasing reward size *w* in each case (Figure 1A-D). This result confirmed our expectation that when reward size is held constant for days and no other water is available, rats will work harder for water when rewards are smaller. An obvious explanation of this could be that each rat performed the number of trials required to earn some fixed amount of water, regardless of reward size (Figure 1A-D, blue curves). We take this as the baseline hypothesis of the study.

**Figure 1.**
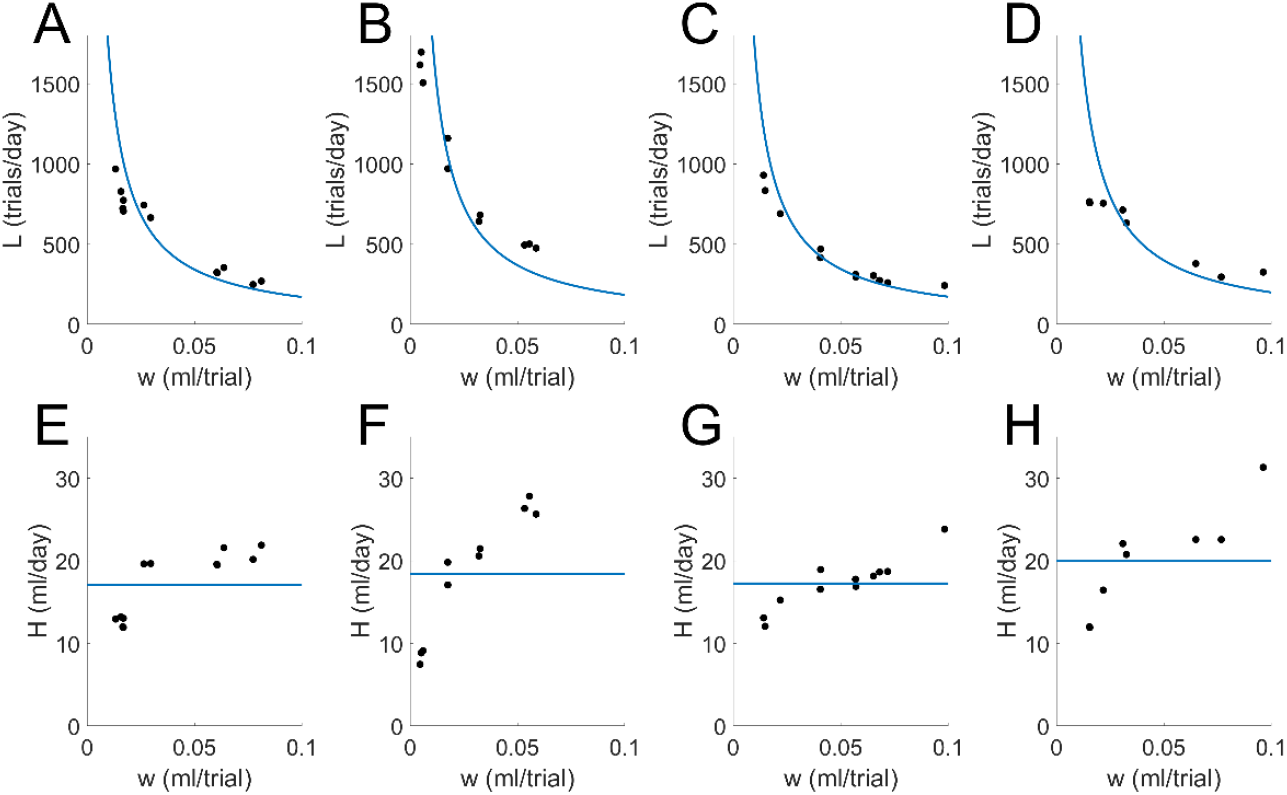
Rats worked harder for smaller rewards, but consumed more water when rewards were larger. Each panel shows results from one rat with 24 hour/day task access. Each symbol shows results averaged over a contiguous stretch of 4 to 7 days on a fixed reward condition, excluding the first day after a reward change. **A-D**: the expected reward *w* (ml/trial) vs. the number of trials performed per day *L*. Blue curves show the best fit to a fixed-intake model *L* = *k*/*w*. **A**. Rat with N=12 steady-state time blocks; the range of reward sizes spanned 0.07 ml/trial; Fixed intake model *k* = 17.1 ml/day. **B**. Rat with N=10 blocks, range 0.05 ml/trial, *k* = 18.4. **C**. Rat with N=11 blocks, range 0.08 ml/trial, *k* = 17.3. **D**. Rat with N=8 blocks, range 0.08 ml/trial, *k* = 20.0. **E-H**: Water intake *H* (ml/day) vs the expected reward *w* (ml/trial), for the same data shown in A-D. Blue lines indicate the best fit fixed-intake model. For statistics see Table 1 in Supplemental Information.

We found, however, that total water intake was positively correlated with reward size (Figure 1E-H; Table 1 in Supplemental Information). This is inconsistent with the fixed-intake baseline model (blue lines), and suggests that the rats took into consideration the cost of water (in effort) when deciding how much to consume. Note that for every reward size shown, the daily water intake was sufficient for the rat to sustain clinically normal hydration, weight, and apparent health and wellness long-term.

### Utility maximization model

We recognized these observations as analogous to phenomena first described in human economics. First, rats did less work per day when rewards were larger, just as in economics the labor supply declines as wage rates increase (*8*). In the case of humans, this so-called back-bending labor supply curve is attributed to workers preferring increased leisure time over increased income. Second, rats consumed more water per day when it was cheaper (in trials/ml), just as in human economics the consumption of a commodity can be sensitive to price, an effect known as price elasticity of demand (*7*). In microeconomics, these patterns have classically been explained by utility maximization theory; therefore, we propose a utility maximization theory to explain our rats’ behavior.

Utility is a measure of subjective value of anything, in arbitrary units of “utils”. Utility can be either positive (for benefits) or negative (for costs). Utility maximization theory posits that individual choices are made to maximize the individual’s net utility, subject to external constraints such as wage rates or prices. The theory allows for utility to be subjective in the sense that preferences can be individually idiosyncratic, but assumes that an individual’s preferences are stable and that individuals are able to determine what behavioral choices will maximize their utility. To apply this theory to our task, we needed to develop a specific utility model for rats doing trials for water rewards.

#### Utility of water

The equation we suggest for the utility of consuming *H* ml of water in a day has only a single free parameter, α:

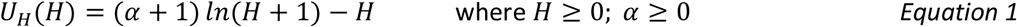

We chose this expression because it met the following criteria: *U*_*H*_(0) = 0 (consuming no water has no utility), it has a positive but decreasing slope (diminishing returns) up to a single maximum, beyond which it declines (consuming too much water is also bad). The maximum of *U*_*H*_(*H*) occurs at *H* = *α*. Therefore *α* has a natural physical interpretation as the rat’s most preferred quantity of water in the absence of any cost. The Marginal Utility of water, the derivative of Utility, is thus:

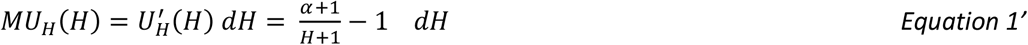

The amount of water consumed *H* is determined by the number of trials done *L* and the reward size or “wage rate” *w*: *H* = *wL* (the *budget line* in economic terms). Thus, we can rewrite Equation 1 in terms of the number of trials done:

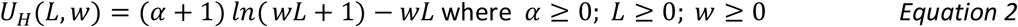

The maximum of *U*_*H*_(*L*) (Figure 2A, Equation 2) occurs at *wL* = *α*. The Marginal Utility with respect to trial number *L* is given by:

**Figure 2.**
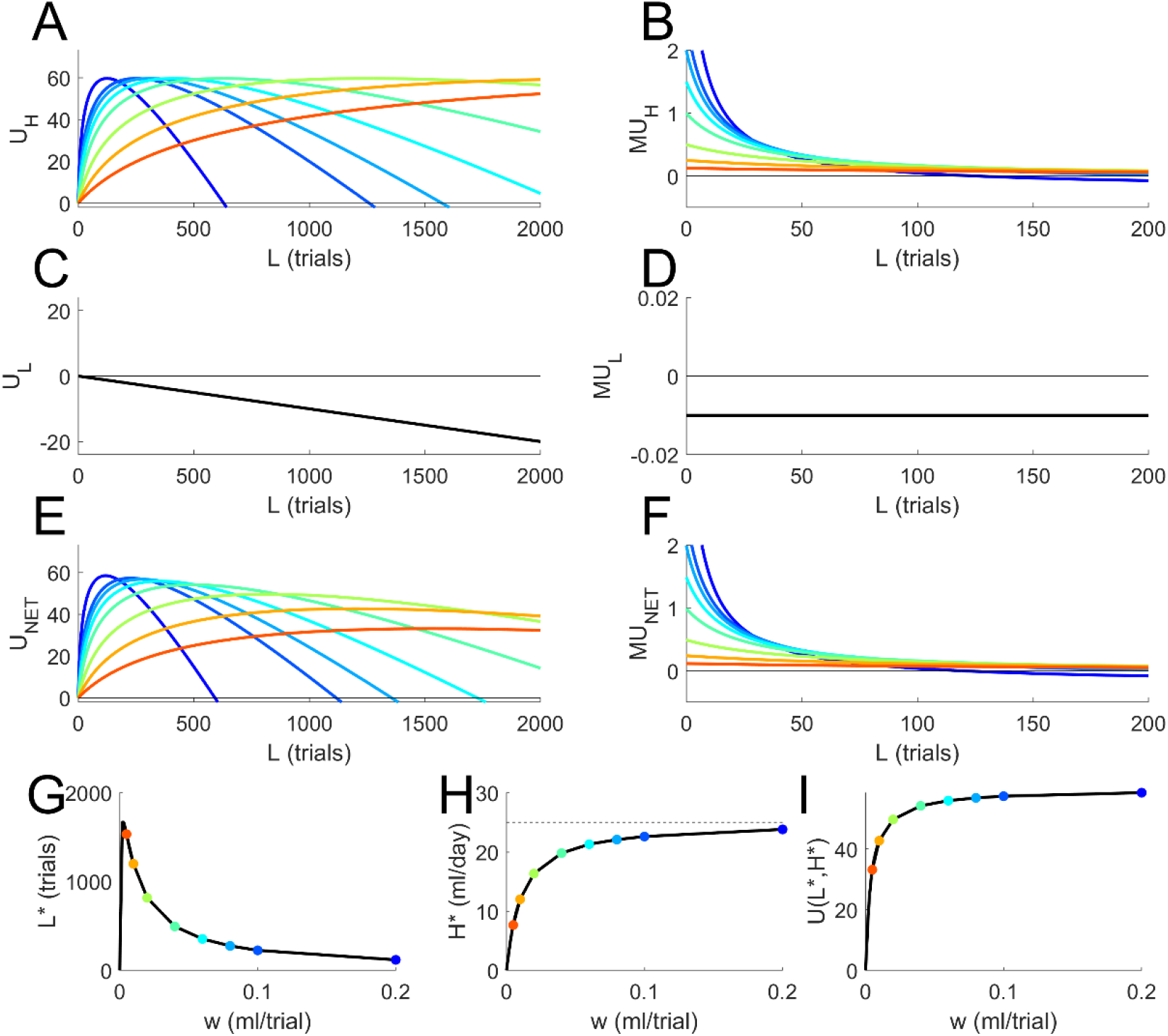
An instantiation of the proposed utility model. Utility model evaluated for parameters α=25, β=0.01. Wage rate (expected reward) is indicated by color, increasing from red to blue: w=0.005, 0.010, 0.020, 0.040, 0.060, 0.080, 0.100, or 0.200 ml/trial. **A**. Utility of water earned by performing *L* trials in one day, in arbitrary units of utils (c.f. Equation 2). **B**. Marginal utility of water with respect to trial number (the derivatives of curves in Panel A; c.f. Equation 2’), evaluated discretely at *dL*=1. Scale is expanded to show detail near origin. **C**. Utility of the work of performing *L* trials in one day, which is negative and does not depend on *w* (c.f. Equation 3). **D**. Marginal utility of labor with respect to trial number (the derivatives of curves in Panel C; c.f. Equation 3’). **E**. Net utility of performing *L* trials in one day, equal to the utility of water (panel A) plus the utility of labor (panel C), c.f. Equation 4. **F**. Net marginal utility of performing *L* trials, the derivatives of the curves in E, or the sum of the curves in B and D, c.f. Equation 4’. Note expanded scale. **G**. The predicted number of trials *L*\* that will maximize utility, as a function of wage rate; compare to data in Figure 1A-D. **H**. The total water income *H*\* earned by *L*\* trials, as a function of wage rate; compare to data in Figure 1E-H. Dashed line indicates the parameter α. **I**. The utility achieved by performing the optimal number of trials, as a function of wage rate.

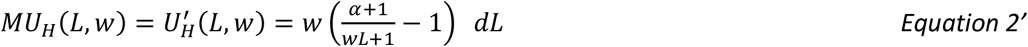

Figure 2A shows curves of *U*_*H*_(*L, w*) for an example value of α and several different wage rates *w*. Figure 2B shows the Marginal Utility curves *MU*_*H*_(*L, w*) vs *L* on an expanded scale near the origin.

#### Disutility of effort

The equation we suggest for the utility of the labor involved in performing *L* trials in a day has only a single free parameter, *β*:

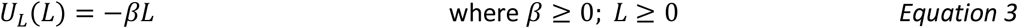

We chose this expression because it was sufficient to meet the following criteria: *U*_*L*_(0) = 0 (doing no work has no cost), otherwise *U*_*L*_ < 0 (working is a cost), and monotonically increasing (more work is more cost). The Marginal Utility with respect to trial number is thus very simply:

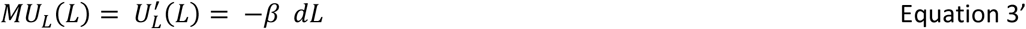

Figure 2C shows the relation of *U*_*L*_ to *L* for an example value of β; Figure 2D illustrates that we are making the approximation that the marginal utility of labor does not depend on *L* in this model.

#### Net utility

Putting these together, the net utility of performing *L* trials to earn *H* water is the sum of the utility of the water and the cost of the effort, as illustrated in Figure 2E.

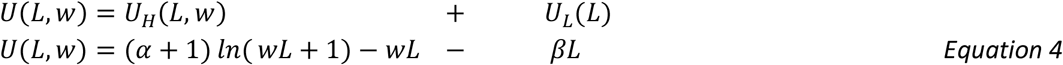

The Marginal Utility with respect to trial number – i.e., the increment in utility of performing *L* + 1 trials compared to *L* trials – is described by the equation (Figure 2F):

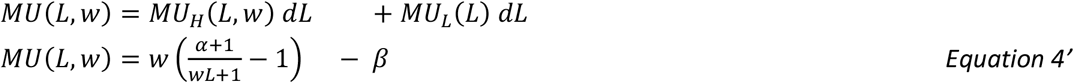

Note that the number of trials done *L* can only be an integer. Therefore, although the functions are defined continuously, we will evaluate them discretely with *dL* = 1.

#### Maximizing utility

In this setting, the utility maximization hypothesis states that rats choose the number of trials that will maximize total utility given the wage rate (the peak of the curve in Figure 2E, or equivalently, the zero-crossing point in Figure 2F). We will denote the optimal number of trials *L*^∗^ and the resulting water intake *H*^∗^.

The null hypothesis that rats simply perform trials until they obtain a fixed target level of water corresponds to the parameter restriction *β* = 0. The maximum utility solutions *L*^∗^(*w*) and *H*^∗^(*w*) are shown in Figure 2G and H respectively for the example parameter choices. Note that as wages get very large, the predicted earned water intake in the task approaches the free water satiety point *α* (Figure 2H, dashed line). The achieved utility *U*(*L*^∗^, *w*) strictly increases with reward size (Figure 2I). This reflects the fact that high-reward environments are always preferable to the rat.

### Fit of the model to the data

We fit the two free parameters of this model to the rat data as described in Methods. Figure 3 panels A-D show the observed trials per day at the granularity of single-day observations (symbols), compared to the maximum utility solution of the model (red curves), for the same rats and experiments shown in Figure 1. The observed (symbols) and predicted (red curves) water consumption as a function of wage rate are shown in panels E-H. We conclude that proposed utility equations are compatible with the qualitative features of example rat data of the type we wish to explain. The fixed income model had the unrealistic implication that as reward size approaches 0, trial rate would approach infinity (panels A-D, blue curves). The utility model predicts that below some minimum wage rate, trial rates fall; and animals will not do any trials if there is no reward (at *w* = 0, *L* = 0).

**Figure 3.**
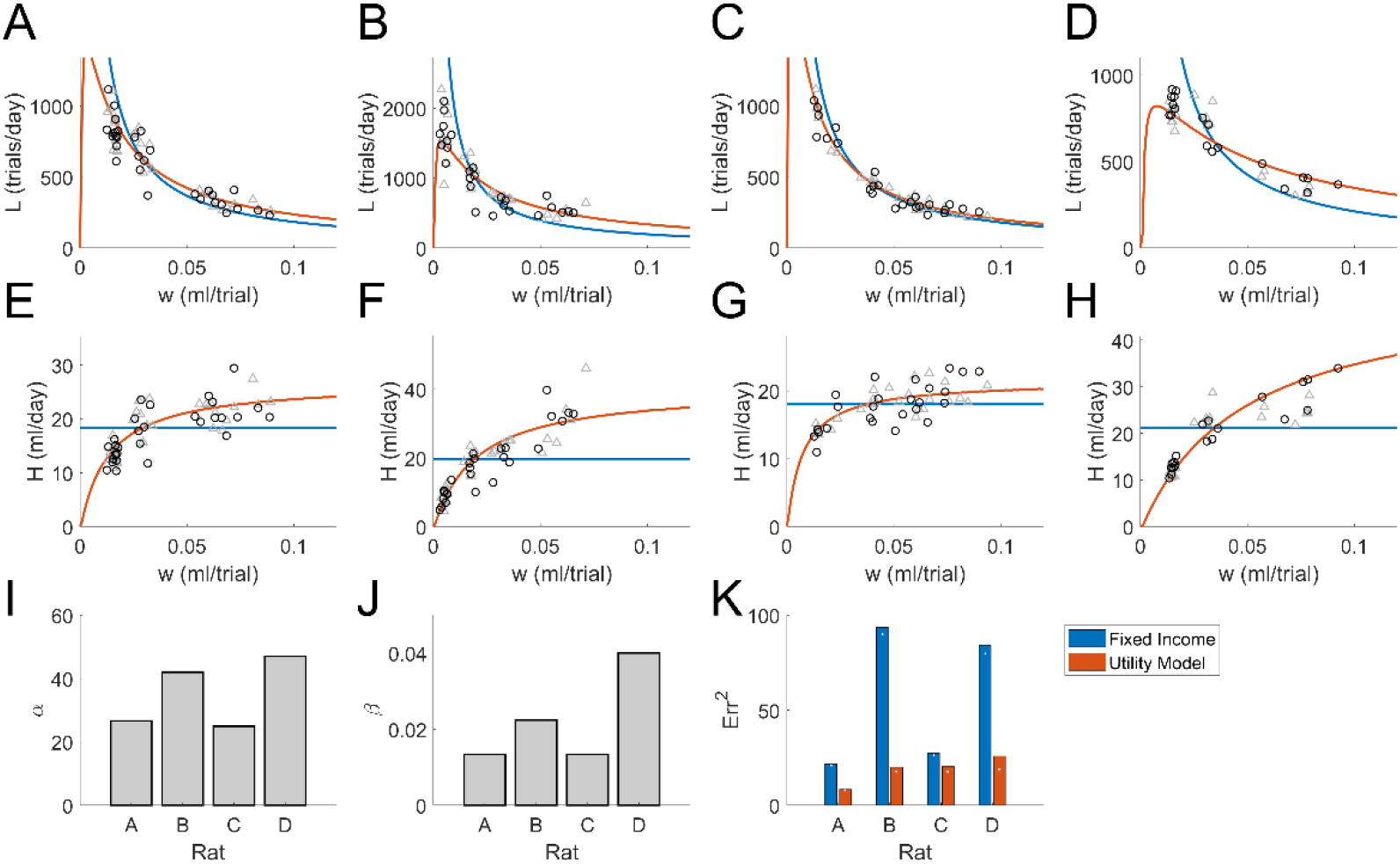
Fit of utility-maximization model to rat data from the 24hr/day task. **A-D**: labor *L* (trials/day) as a function of wage *w* (ml/trial) for the same experiments as Figure 1A-D, shown at single-day resolution (symbols), compared with the utility maximization model (red curve) or fixed income model (blue curve). Half the data points (gray triangles) were used to fit model parameters for the curves shown; black circles show hold-out data. **E-H**: income *H* (ml/day) as a function of wage *w*, for same data and utility model solutions as A-D. **I**. Values of the parameter *α* fit to each rat, averaged over all leave-one-out fits. **J**. Values of the parameter *β*. **K**. The cross-validated residual error of the fixed-income model (blue lines in E-H) and of the utility model (red curves) with respect to water income *H*, based on leave-one-out cross-validation. White points indicate the mean residual error on the fitted data for comparison.

The fit parameter values are shown in Figure 3I-J, and the cross-validated residual error of the fit is compared with that of the fixed-income model in Figure 3K. The parameter *α* predicts the rat’s ad lib water satiety point. Although satiety was not measured in these rats, the values of *α* are plausible 24-hour ad lib water consumption values based on measurements in other adult rats (*12*), Supplemental Figure.

### Access schedule affects effort and consumption

Four rats were tested on both 24 hour/day and 2 hour/day schedules, with a range of reward sizes on at least one schedule. In time-limited sessions, access to the task was limited to two hours/day and no water was given between sessions, such that the water earned in the task was still the rat’s total water intake for the day. Rats can perform trials as rapidly as every four seconds, so in principle a rat could complete up to 1800 trials in a 2-hour session. But the rats did fewer trials and therefore consumed less water per day on the 2-hour condition than the 24-hour condition with a comparable wage rate (Figure 4A-H symbols), as we have noted previously (*12*).

**Figure 4.**
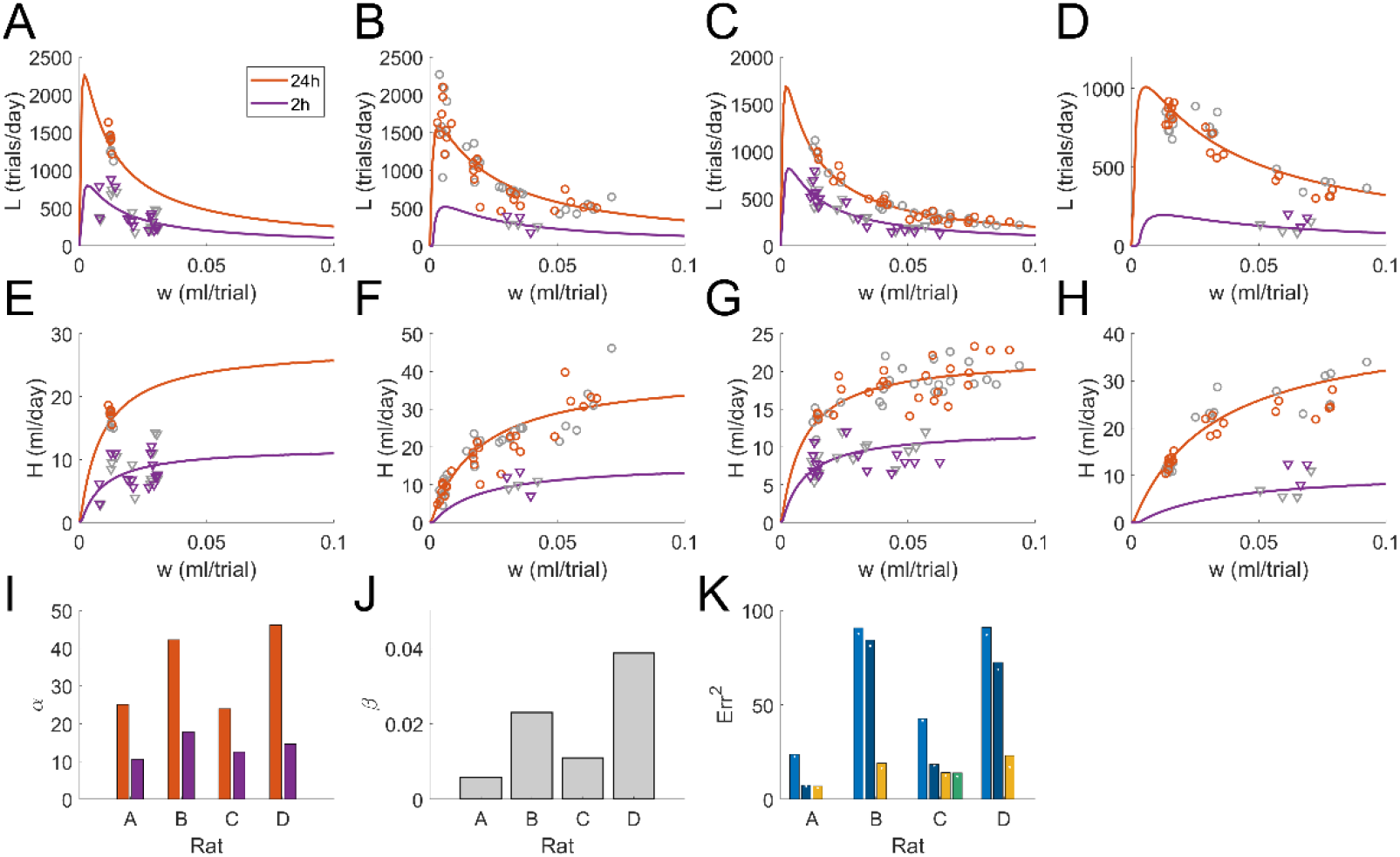
Data and model fits for rats tested on two access schedules. Each symbol represents data from a single day during a steady state period, on either 24 hour/day (red, circles) or 2 hour/day (purple, triangles) access schedules. Curves represent the model fit to both schedule conditions simultaneously, with distinct α parameters and a shared β parameter. Gray symbols indicate data used to fit parameters for the shown curves; colored symbols are hold-out data. Panels A-D: observed effort *L* (symbols) and utility maximizing effort *L*\* (curves). Panels E-H: observed water consumption *H* (symbols) and utility maximizing consumption *H*\* (curves). Panels B-D and F-H are from the same rats as corresponding panels in Figures 1-2. Data in panels A, E are from a different rat not shown above. **I**. The fit values of *α* for the 24hr/day (red) or 2hr/day (purple) conditions, averaged over all leave-one-out fits. **J**. The fit values of *β* (shared by both schedule conditions). **K**. Residual errors by leave-one-out cross-validation, for the best-fit simple fixed income model (fixed target volume regardless of condition, 1 parameter), light blue; the schedule-dependent fixed income model (different fixed target for each schedule, 2 parameters), dark blue; the utility maximization model with schedule-specific α and common β (3 parameters), yellow; or the utility model with schedule-dependent β (4 parameters), in green. Only one had a broad enough range of wage rates on both schedules to fit the four-parameter model. The average residual errors on fitted data are indicated by white points.

Access schedule is known to have a strong effect on daily free water consumption of mice (*13*) and we have observed this in rats as well (Supplemental Figure). This implies that the parameter *α*, which is equal to the free-water satiety point, must depend on the access schedule. To explore whether this effect alone could explain the effect of task scheduling on trials, we fit the utility equation to the data from both schedules jointly, with a distinct satiety parameter *α* for each access schedule (*α*_24_, *α*_2_) and a shared parameter *β*. This approach was surprisingly successful at capturing the main structure in the data (Figure 4A-H curves; 4K).

The free water satiety was not measured for the rats in this study, but the alpha parameters of these fits (Figure 4I) are consistent with our observation in other rats that free water satiety point is both lower and less variable with 2 hour/day access than with 24 hour/day access (Supplemental Figure). Two of these rats were much better explained by utility maximization than the fixed-income model with a different fixed-income target for each schedule (Figure 4K, rat B and D). The other two rats either had less elasticity of demand, and/or their elasticity was only expressed outside the range of reward sizes we tested, and therefore were fit about equally by both models.

The utility equations we propose are based on the data presented; the data should not be taken as a test of the theory. We note that the model does not require that all rats necessarily show elasticity of demand (i.e. the parameter *β* can be 0 for some rats). New rats, including male rats, will need to be tested to determine the distribution of elasticity of demand among rats in general.

### Dynamic interpretation of marginal utility

The utility maximization theory presented above is a static equilibrium model based on maximizing the total utility of doing *L* trials and harvesting *H* water in one day (given the wage rate, and a schedule-dependent satiety parameter *α*). We hypothesize, however, that the mechanism by which rats solve the utility maximization problem is temporally local: the rat continuously estimates the change in utility it would experience for doing *one more* trial, and initiates another trial if and only if the expected change in utility is positive. More strongly, we propose that the probability of doing a trial at any moment is a monotonic increasing function of the net marginal utility with respect to trial number (Equation 4’). The trial rate cannot be less than zero and has a physical upper limit of ∼0.25 trials/sec. Therefore, we expect a sigmoid relationship between the time-varying estimate *MU*(*t*) and the trial rate.

We explored temporal dynamics in 2-hour sessions because these are phase aligned by the start time of the sessions, and limited to a short fraction of a circadian cycle. We fit the model parameters using the per-day trial counts for all schedules and wage rates (Figure 4) and used the parameters to generate marginal utility curves for the wage rate of interest in each case (Figure 5A-B).

**Figure 5.**
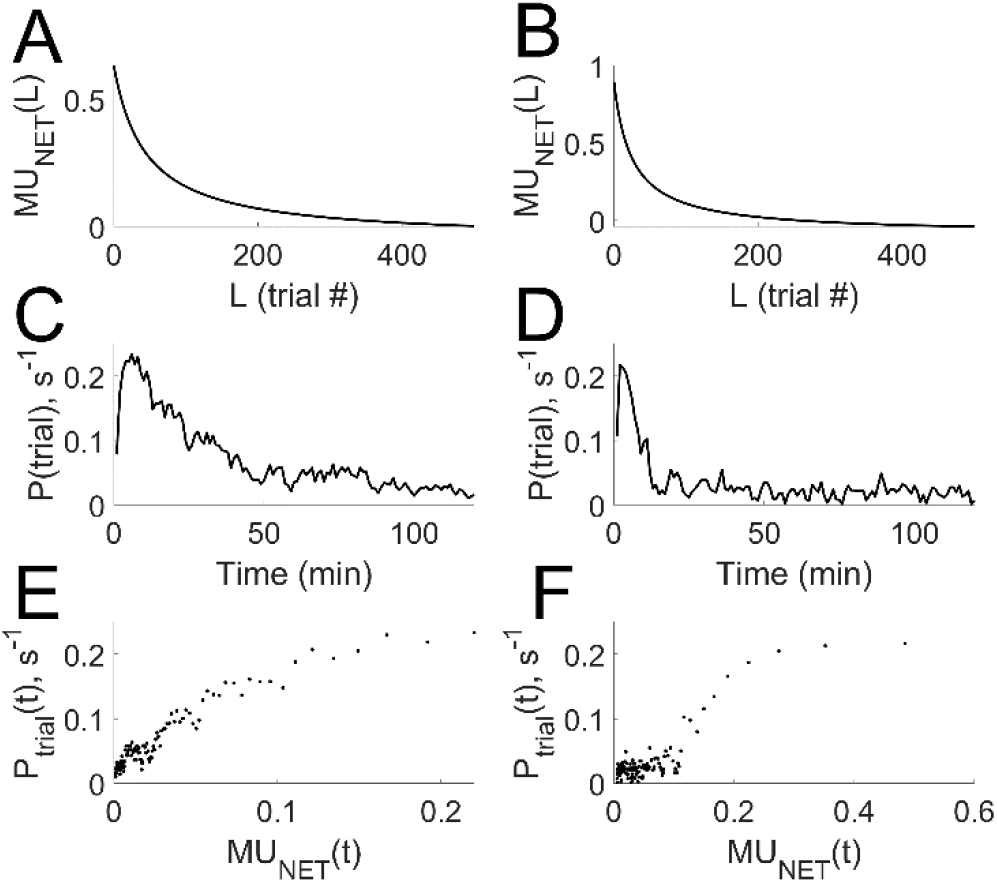
Reinterpretation of the equilibrium model as a time-varying function. **A**. Marginal Utility as a function of trial number (Equation 4’) for the 2hr/day schedule, based on the parameters (*α*_2_ = 29.5, *β* = 0.031) fit to the rat’s daily trial counts on both schedules and all wage rates (Figure 4), and evaluated for *w* = 0.023. **B**. Like panel A for a different rat and wage rate. Parameters *α*_2_ = 23.5, *β* = 0.046, evaluated for *w* = 0.040. **C**. Observed trial density over time in N=33 two-hour daily sessions with wage rate *w* = 0.023 +/−0.002, for the rat whose *MU*(*L*) curve is shown in A. **D**. Like C for the rat whose *MU*(*L*) curve is shown in B, N=14 two-hour daily sessions with *w* = 0.040 +/−0.003. **E**. The marginal utility at each time point (*MU*(*t*), determined by *MU*(*L*) for the average cumulative number of trials *L* at time *t*) is compared with the observed instantaneous rate of trial initiation, for the case analyzed in A and C. **F**. Like E, for the case analyzed in B and D. Examples were chosen as the two cases in which the same wage rate was tested for the most consecutive days in the 2-hour schedule. Different rats from any shown above.

If *MU*(*L*) is the instantaneous drive to initiate a trial after the *L*_*th*_ trial, the rats’ trial rates should drop steeply during a session: *MU*(*L*) fell to half its initial value by trial 39 (Figure 5A) or by trial 22 (Figure 5B), in both cases corresponding to the rat having consumed only 0.9 ml of water after 22 hours of water restriction. The observed timing of trials in these sessions were qualitatively consistent with this prediction (Figure 5 C-D). The probability of initiating a trial at time *t* is a sigmoidal function of marginal utility (Figure 5 E,F).

### Quantitative predictions

A strength of the utility model is that it can be fit with very few free parameters and makes several quantitative predictions, including experimental manipulations not used in derivation of the model. First, the parameter *α* represents the rat’s free water satiety point. Therefore, *α* could be constrained by a free water satiety measurement, leaving only one free parameter *β* to explain daily trial number and water consumption as a function of reward size on a given schedule (e.g., Figure 6A, black curve).

**Figure 6.**
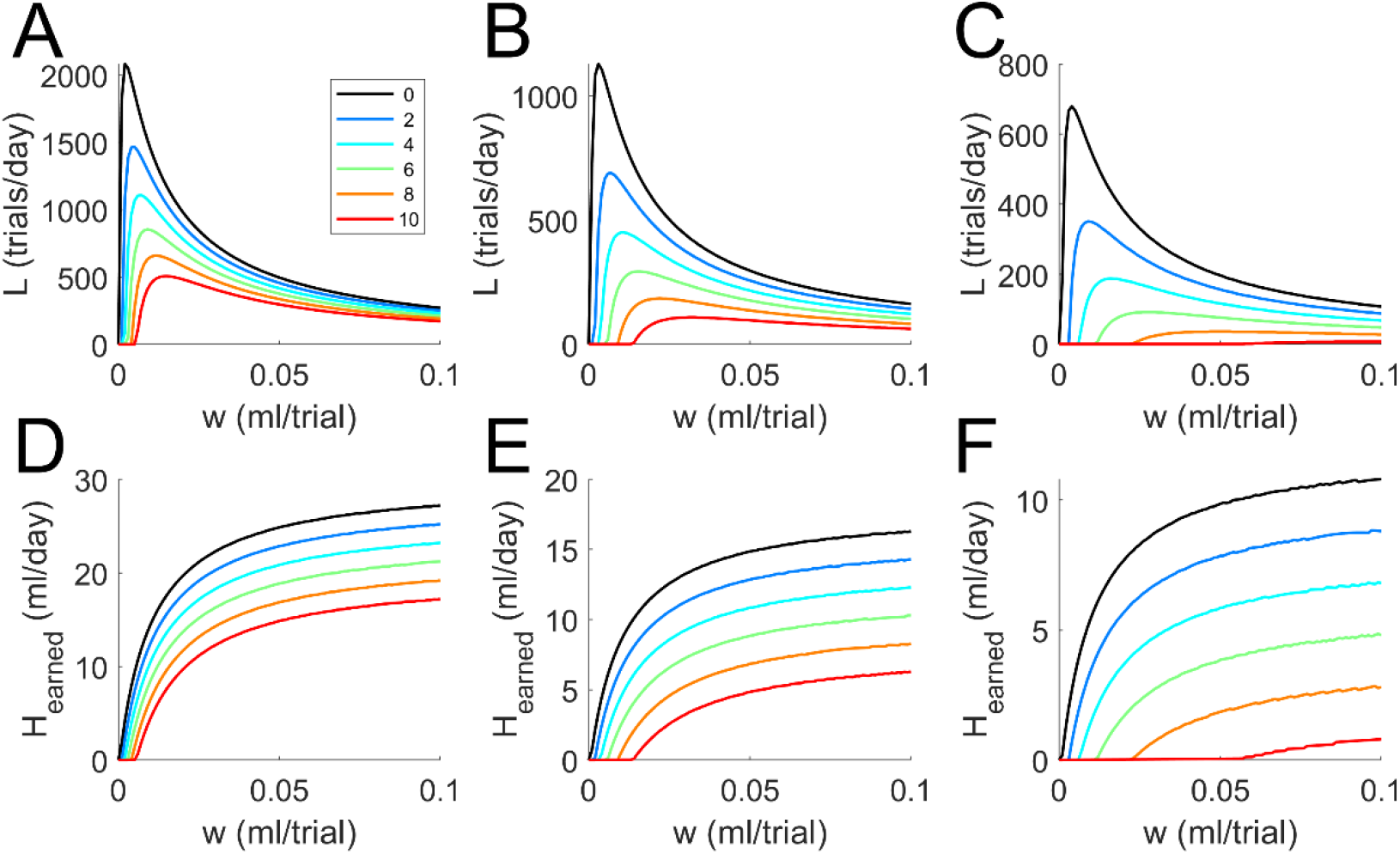
Quantitative predictions of the model. **A**. Utility-maximizing trial rate *L* as a function of reward size *w*. The single free parameter *β* for a rat could be fit using observed daily trial counts from a range of reward sizes *w* tested on a 24 hour/day schedule with no endowment, and experimentally measured 24-hour free water consumption (here hypothetically *α*_24_ = 30 ml/day), producing the model curve shown in black. Without additional free parameters, the model predicts the trial rate for any reward size in the presence of any free water endowment *H*_0_ (ml/day, color key). **B-C:** With measured free water consumption on two other schedules, here hypothetically 8 hours/day *α*_8_ = 18 ml/day (B) and 2 hours/day *α*_2_ = 12 ml/day (C), the model further predicts the trial number for any novel combination of schedule, endowment, and reward size with no additional free parameters. **D-F:** The earned income *H*_*earned*_ = *wL* corresponding to the trial numbers predicted in A-C. Note that the rat’s total water intake, not shown, includes the endowment (*H*_*total*_ = *H*_0_ + *wL*).

Second, after fitting the parameter *β*, the model makes a quantitative prediction for the effect of a supplement or “endowment” of daily free water (Figure 6A, colored curves) with no additional free parameters. This is nontrivial because endowments shift the utility of water curve (Figure 2A) horizontally relative to the utility of labor curve (Figure 2C), resulting in changes in trial rate and total income that depend nonlinearly on the wage rate.

Third, the model predicts a forward-bending part of the labor supply curve (*L* initially increases with *w*). In the present study, ethical considerations prevented us from testing reward sizes that would have resulted in insufficient water intake. In the proposed endowment experiment, however, the forward phase of the labor curve sometimes overlaps with conditions that provide adequate daily fluids, so it should be observable.

Fourth, we suggest that the effect of access schedule on water satiety may be sufficient to account for the effect of schedule on labor (c.f. Figure 4). If so, one could fit the parameter *β* using data from one schedule (black curve in Figure 6A), and use this to predict the trial rate for any other combination of schedule, wage rate, and endowment (i.e. all the other curves in all the panels of Figure 6) with no additional free parameters, constraining *α* by measured free water satiety on each schedule. This could be compared to the alternative possibility that *β* is also schedule dependent.

In summary, the proposed theory has the potential to quantitatively explain the nonlinear interacting effects of three environmental variables (wage rate, schedule, and endowment) on rats’ willingness to work for water, with as few as one free parameter (when *α* is empirically constrained, if *β* proves to be schedule-independent), or at most one free parameter per schedule (if *β* depends on schedule). Any deviations from predictions will be informative for revising the analytic form of the model, which would alter the shape of the utility curves and therefore update the predictions for neural dynamics. Alternative utility equations are considered in Supplemental Information.

### A neural hypothesis

We have proposed that marginal utility *MU* could be re-interpreted dynamically, and showed that this is consistent with the timing of behavior (Figure 5). On this hypothesis, the rat’s task of solving the utility maximization problem reduces to simply detecting whether *MU* > 0 at any given moment. This raises the question of where in the brain *MU* is computed. The recent explosion of progress in unravelling the neurobiology of thirst (*14-30*) provides an unprecedented opportunity to link behavioral motivation to known neural mechanisms within an economic theory framework.

The subfornical organ (SFO) is part of *lamina terminalis*, a circumventricular forebrain nucleus involved in the regulation of thirst. Within *lamina terminalis*, SFO is tightly interconnected with the median preoptic nucleus (MnPO) and the organum vasculosum of the lamina terminalis (OVLT). Beyond *lamina terminalis*, SFO projects to the paraventricular nucleus and the supraoptic nucleus (SON) of the hypothalamus.

The glutamatergic neurons in SFO directly sense plasma osmolality, as well as integrating other signals of physiological hydration (*16*). Activity in SFO^GLUT^ neurons is high in dehydrated animals, and declines rapidly as soon as water is ingested, long before physiological hydration is restored (*16, 18, 20, 29*). If activity in these neurons is artificially suppressed, dehydrated animals will not drink (*18*). If activity is artificially induced, water-sated animals drink voraciously (*18, 21-23*). These observations established a central causal role of SFO^GLUT^ in regulation of water ingestion, and mirror the properties expected of the neurons that compute marginal utility in our task.

The time course of *MU*(*t*) after the onset of task availability in rats (Figure 7A) resembles the time course of activity of SFO^GLUT^ neurons in the first minutes after thirsty mice access water (Figure 7D). The steep decline in the probability of rats initiating trials (Figure 7B) resembles the steep decline in the probability of the mice licking a water tube (Figure 7E). Utility theory provides a functional explanation for why SFO^GLUT^ neurons stop firing and mice stop drinking long before they are physiologically hydrated: the neurons do not represent the hydration state of the animal, rather they represent the expected marginal utility of additional consumption.

**Figure 7.**
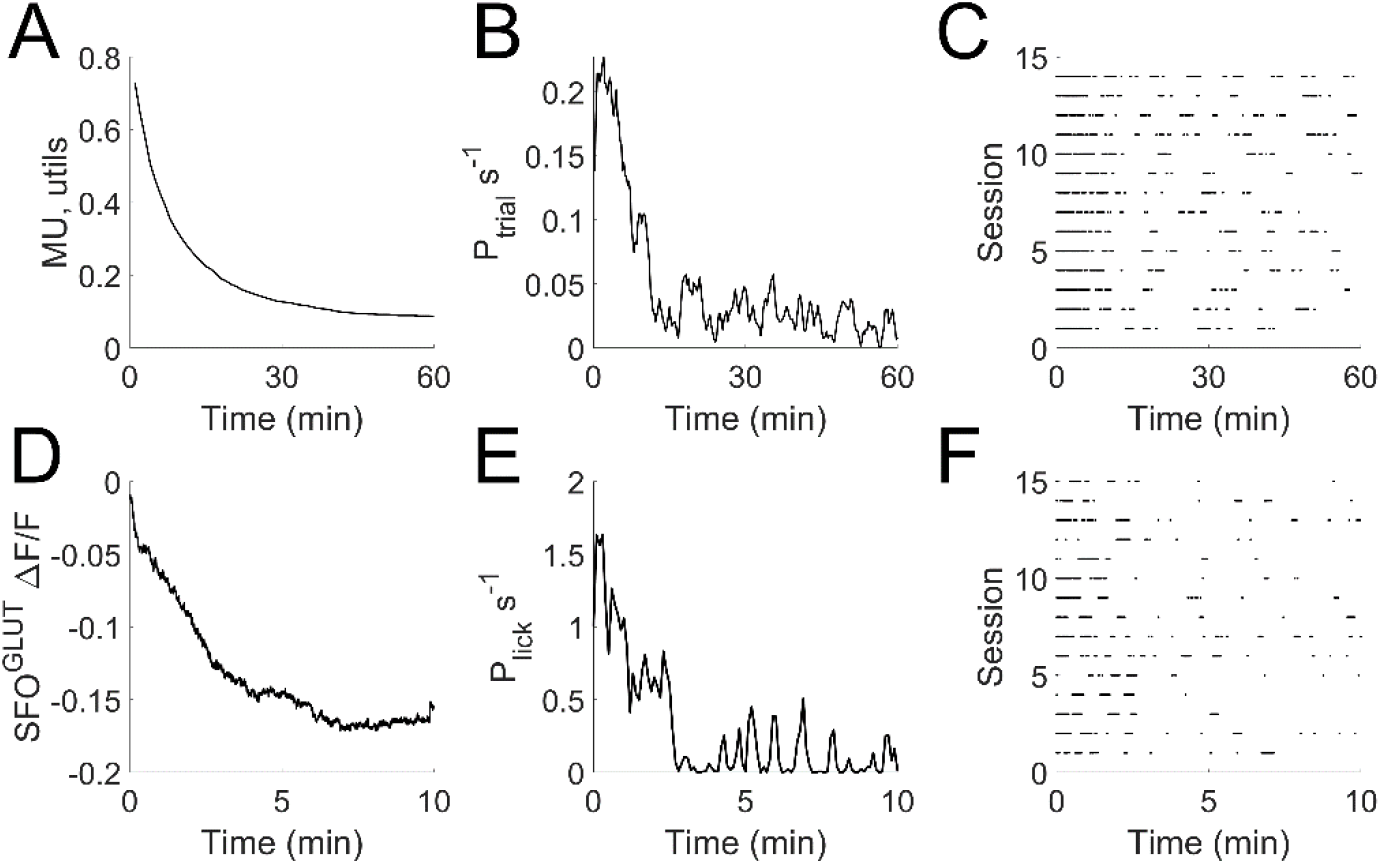
Hypothesis: SFO^GLUT^ activity is the neural representation of MU. **A**. Marginal Utility expressed as a function of time, based on the parameters fit to one rat’s daily trial count as a function of wage rate and access schedule, c.f. Figure 5B,D,E. **B**. Observed trial rate as a function of time in 14 consecutive 2-hour sessions by one rat tested on a 2 hour/day schedule at the wage rate modeled in A (c.f. Figure 5D). **C**. Times of trial initiation for the first hour of each 2-hour session (rows), each point indicates the time of one trial. **Panels D-F:** data from mice at the onset of water access after water restriction, from (*29*) and additional unpublished data from that study. **D**. Activity of genetically identified SFO^GLUT^ neurons at onset of drinking, measured by fiber photometry. Population activity is expressed as the change in GCaMP fluorescence (%) relative to preceding baseline, averaged over N=15 sessions in 15 different mice. **E**. Average licking rate in first 10 minutes after water access from the same experiments as D. **F**. Times of licks, where each row is an individual session and each lick is indicated by a point.

The rapid shutoff of SFO^GLUT^ activity upon drinking is mediated by at least two sensory feedback mechanisms. One depends on proprioceptive sensations of swallowing and involves feedback to SFO from inhibitory neurons in MnPO (*14, 18*). A second mechanism involves vagal feedback from osmotic sensors in the gut (*29*). These circuits provide candidate neural mechanisms for the how SFO rapidly updates expected marginal utility. On average, SFO^GLUT^ activity falls off smoothly with time during drinking, but in any individual session SFO^GLUT^ activity oscillates around this average (*29*), such that the licks occur in bursts (Figure 7F). We observe similar bursts in the timing of trials in individual rat sessions (Figure 7C), consistent with the hypothesis that *MU* is computed by SFO^GLUT^ in rats.

A direct test of this hypothesis would require recording from and causally manipulating these neurons during task performance. But known properties of SFO^GLUT^ neurons allow some predictions that could be tested in much simpler behavioral experiments. First, non-hydrating fluids such as hypertonic saline or oil are sufficient to trigger the rapid inhibitory feedback signals to SFO^GLUT^ that underlie the anticipatory or predictive drop in drinking behavior (*18, 20*). In our task, probe sessions using non-hydrating fluid rewards instead of water rewards should therefore exhibit the same rapid drop in *MU* and in trial rate at early times in short sessions, despite the lack of any relief of the rat’s dehydration. Conversely, hydrating fluids that bypass both oral and gut sensory neurons should fail to trigger these rapid inhibitory feedback circuits to SFO^GLUT^(*18, 20*). In our task, therefore, if MU is computed by SFO^GLUT^ neurons, free water endowments provided as subcutaneous or IV fluids just before a short session should have much less effect on the trial rate dynamics or total trials than the same endowment given orally – in spite of more rapidly relieving the rat’s dehydration.

## Discussion

The affordances of the environment change over time. Therefore, animals must regulate their allocation of effort flexibly to ensure that each basic need is met without wasting time and energy. In our experiment, rats did less work per day to earn water during protracted periods when water rewards were larger. Moreover the rats consumed substantially more water per day when it was easier to earn, consistent with elasticity of demand in economic theory (*7*).

These findings are in line with classic animal behavior studies showing that animals behave “rationally” in the sense required by utility maximization theory (*31-42*). Our study extends the literature in several respects. The “work” in our task included components of mental effort (difficult perceptual discriminations), reward uncertainty (the probability of reward in a single trial was <1), and risk of punishment (time-outs on unrewarded trials), as opposed to merely mechanical effort (lever presses) studied previously. The reward in this study was water, rather than food or food-and-water compound rewards used previously. The classic experiments compared three wage conditions per experiment (baseline, uncompensated wage change, and compensated wage change), and assessed only a qualitative outcome (e.g. the direction of substitution and income effects being compatible or incompatible with the theory). To our knowledge none of the past studies explored the large number of different uncompensated wage changes sufficient to define the shape of the utility curves as we do here. Unlike classic studies our experiments recorded the time of every unit of labor and unit of water consumption, allowing us to relate MU to the temporal dynamics of behavior. Other recent studies have also fruitfully revisited and extended animal models of other aspects of economic theory (*43-46*).

Our results can be reconciled with an apparently contradictory recent study in rodents (*3*), which showed that the vigor of response to a single available option is positively related to reward size. In that study, reward size was changed many times within a single 2-hour session. On this short timescale, fatigue from harvesting small rewards could reduce the animal’s ability to harvest when rewards are larger; and animals can easily survive without any water for a few minutes while waiting for a better opportunity. In this context, the strategy of working harder for larger rewards pays off. In other words, rapid successive options are more comparable to simultaneous value comparisons. The previous study was also designed to equalize the state of thirst or satiety across compared reward conditions. Our experiments are complementary to these, in that we explored reward changes on much longer timescales, and in the presence of an intact feedback loop whereby behavioral choices had consequences for internal states that directly altered subsequent motivations. The fact that rats adopt different strategies in the two experiments implies that rats learn not only the expected volume of the next water reward, but also the temporal correlation of reward volume changes. How and where these estimates are computed remains unknown.

Variability in trial rate may reflect uncertainty in the rat’s estimation of *MU*(*L*). In a closed economy, a consistent misestimation of *MU*(*L*) would result in cumulative physiological dehydration or overhydration, which would provide a corrective feedback signal to the utility estimate. Some of the unexplained variance may be attributable to biological rhythms. Although we analyzed data at one-day granularity, the timing of circadian bouts of activity could vary relative to the time of experimental observations, causing day to day variability. Female rats tend to eat and drink less during estrous, which could cause variation on a 4-day cycle. Ambient temperature and light cycle were held constant throughout the year, but we cannot rule out seasonal fluctuations. Finally, the data were collected over months to years, so age effects could also contribute to variability.

Why would a rat consume more water than it needs? The capacity to store water in body tissue is very limited; unused intake is soon eliminated as urine. Yet rats that were able to maintain good health on 8-10 ml/day with small rewards were willing to do work to get three times that much water when rewards were large. This aligns with our previous finding that if water is rendered unpalatable, rats consume about 10 ml/day and maintain health, but will consume 20-40 ml/day when water is plain (*12*). We speculate that rats use this optional extra water in part to enable extra dry food consumption; unlike water, excess calories can be stored. This could be tested by measuring or restricting food consumption. Other uses of excess water might include increased exercise/exploration, increased grooming, or reducing the amount of energy required for fluid retention or waste elimination in the kidney.

The utility maximization framework could be applied to tasks with different rewards (such as food) or costs (such as predation risk, time investment, or caloric expenditure). Each good or cost would have its own characteristic utility equation and underlying neural computations. When multiple goods and costs are involved, interactions among them can be stipulated as constraints limiting the available options, and the component marginal utilities added together to compute net marginal utility of each option.

We have advanced the hypothesis that net marginal utility ***MU***_***NET***_ (Figure 2F) is computed by the glutamatergic neurons SFO^GLUT^. These neurons drive glutamatergic neurons in the median preoptic nucleus (MnPO^GLUT^), which are also causally required for drinking behavior (*14*). A variant of this hypothesis is that SFO^GLUT^ neurons compute the marginal utility of water ***MU***_***H***_ (Figure 2B) and this is combined with other information to compute net marginal utility downstream, either in MnPO^GLUT^ or later. Dopamine circuits have long been implicated in response vigor and willingness to work (*47-49*), as well as in representation of marginal utility (*50*). The SFO^GLUT^ neurons modulate phasic responses in dopamine circuits during drinking (*28*), so dopamine circuits that encode ***MU*** in this task could inherit this information from SFO. Signals in insular and cingulate cortex related to predicted water need or hedonic water value (*24-27*) may also be downstream of computation of ***MU*** in SFO.

### Summary

We have presented experimental data showing that when rats had to work to earn their water, they worked harder for smaller rewards, but worked for more total water when it was easier to get (Figure 1). We propose an analytic utility-maximization model (Figure 2) that is able to account for these observations (Figure 3) and that suggests an explanation for the effect of access schedule (Figure 4). We suggest a dynamic re-interpretation of marginal utility and relate this to the observed timing of behavior (Figure 5). The model makes testable quantitative predictions, with the potential to explain the dependence of behavior on three environmental variables (wage rate, schedule, and endowment) with one or a few free parameters (Figure 6). We advance the new hypothesis that SFO^GLUT^ neurons compute the model variable *MU* (Figure 7) and suggest how this can be tested. The model thus spans descriptive, quantitative, normative, algorithmic and mechanistic levels of explanation.

## Methods

### Experimental

All experiments were performed in strict accordance with all international, federal and local regulations and guidelines for animal welfare. Experiments were performed in AAALAC accredited facilities with the approval and under the supervision of the Institutional Animal Care and Use Committee at the University of California, San Diego (IACUC protocol #S04135).

The eligible cohort for this study contained 16 female Long-Evans rats (Harlan Laboratories, Indianapolis, IN). Of these, ten had been previously tested for effects of ad lib citric acid water on trial rate (*12*) and were older adults, and six were naïve young adult rats. Males remain to be tested in a future study. Although the sample size was too small to test effects of age, water satiety point appeared to be higher in the older adult females compared to the young adult females, suggesting that satiety should be tested as a function of age in long-duration experiments. Additional details in Supplemental Information.

The task the rats performed for work was a random dot motion visual discrimination task (*10*) conducted using a custom automated training and testing system (*11, 51*) whose control software is written in MATLAB (MathWorks, Natick MA). Briefly, an operant chamber was connected to the animal’s home cage by a tube either chronically (for the 24 hour/day schedule) or for a daily timed session (for the 2 hour/day schedule). In the operant chamber there were three infrared beam-break lick sensors arrayed along the bottom edge of an LCD monitor visual display. The rat was required to lick the sensor at the horizontal center of the screen to initiate a trial, at which time the visual motion stimulus appeared and persisted until the rat licked a response sensor on the right or left side. A response lick on the side toward which the visual motion flowed was rewarded with a drop of water. Incorrect responses were punished by a 2-second time-out. The rewarded side was selected randomly with equal probability independently each trial. Because the visual motion signal was embedded in noise, rats made errors and thus received rewards in approximately 75% of trials. Rats were individually caged during task access, but pair-housed between timed sessions for the 2hour/day condition. Dry rat chow was continuously available during and between sessions. The shaping sequence to train rats to perform the task has been described elsewhere (*10, 11*).

Within the task software, reward volume was controlled by the duration a solenoid valve opened to allow flow from a gravity-fed water source (a 30-to 60-ml calibrated syringe filled to standard level daily and positioned at a fixed height above the chamber). This nominal reward size (valve open time) was held constant for at least four and up to 14 consecutive days. Rats received daily health checks and were removed from the experiment immediately if they experienced >10% weight loss or showed clinical signs of dehydration. Therefore, we only report results for reward sizes for which a rat was able to maintain body weight and clinically normal hydration for at least four consecutive days without water supplements. Additional details in Supplemental Information.

### Analysis

Analysis and modeling were performed in MATLAB version R2018b. The trial data were automatically recorded by task software, body weights and water consumption data were manually entered at the time of daily observations. These data were later aggregated by scripts which identified all consecutive stretches of dates with constant expected reward (ml/trial) and access schedule (∼24 hours/day or ∼2 hours/day), and no free water supplements. The first date after a change in either schedule or reward size was excluded from analysis to allow for possible transition effects. Any missing or inconsistent data points were resolved by inspection of written lab notebooks. No missing data were simulated or interpolated. The effective wage rate was estimated from the measured water consumption and observed trial number on a daily basis whenever direct measurements were available, or inferred from calibration otherwise, as detailed in Supplemental Information.

To find the parameter combination for the utility model that minimized the mean squared error of prediction, we performed an exhaustive progressive grid search. The error surface was convex and the minimum squared error solution unique. To find the maximum utility solutions for a given parameter combination, Equation 4 was numerically evaluated for all integer values *L* from 0 to 10^5^, at each experimentally tested wage rate *w*. For model selection purposes we estimated residual errors by leave-one-out cross-validation, predicting each data point with a model that was fit to the other data points (Figures 3K and 4K). Additional details in Supplemental Information.

### SFO Data

Calcium imaging data from the subfornical organ (SFO) in mice during water consumption (Figure 7D-F) were from a previous study (*29*). For detailed methods see (*18, 29*). Briefly, a recombinant AAV expressing the fluorescent calcium indicator GCaMP6s was injected into the SFO of Nos1-IRES-Cre mice, resulting in expression in a genetically defined population of glutamatergic neurons. A fiberoptic cannula was implanted above the SFO for fiber photometry. The imaged calcium fluorescence was normalized using the function: ΔF/F = (F – F_0_)/F_0_, where F_0_ is the median fluorescence of the baseline period prior to water access.

## Data and Code Availability Statement

An interactive, executable replication code and data repository is available at CodeOcean (*52*). This repository includes documentation of intermediate analyses and contains all data not shown.

## Acknowledgements

Nicole Dones made the initial observation prompting this study, and assisted in data collection and curation. Neehar Kondapaneni assisted in data collection and curation and piloted earlier models. Carly Shevinsky provided expert technical assistance. Daily health monitoring of animals was also provided by Serena Park, Ryan Makin, Anjali Herekar, Michaela Juels, Xiao Guo, Vishal Venkatraman, and Grace Lo. Christopher Zimmerman and Zachary Knight offered valuable discussions about thirst neurobiology, and shared the data from mouse SFO used in Figure 7. Blake Bruell helped develop Equation 4. Mark Machina provided helpful discussions about economic theory and commented on early drafts of the manuscript. Support for this work was provided by a Regents of California Academic Senate Grant R0034B.

