## Supplemental Information for "*Rattus economicus*: A neural model for rational regulation of water-seeking effort in rodents"

Pamela Reinagel

Division of Biological Sciences, Neurobiology Section, University of California San Diego, La Jolla CA 92103.

#### Symbol Definitions

| Variable | Name | Units | Bounds | Description |
|---|---|---|---|---|
| w | Wage rate | ml/trial | ≥ 0 | Expected reward (ml/reward x reward/trial) |
| L | Labor | trials/day | ≥ 0 | Total trials performed by rat per 24-hour period |
| $H_0$ | Endowment | ml/day | ≥ 0 | Free water given ($H_0$=0 in all experiments shown) |
| H | Income | ml/day | ≥ 0 | Total water consumed by rat per 24-hour period, $H=H_0+wL$ (H=wL in all experiments shown) |
| $\alpha_x$ | alpha, for schedule x | ml/day (hours/day) | > 0 | Parameter for utility of water (schedule-specific); equal to the free-water satiety point for schedule. |
| β | beta | unitless | ≥ 0 | Parameter for disutility of effort |

#### Supplemental Methods

##### Sampling Details

Data were accrued up to a pre-scheduled end date, except that some older adults were removed from the study sooner due to age-related disease. The number of time blocks or distinct reward sizes after data curation differed between rats for many reasons: some but not all of rats had qualifying data that were collected in the course of the previous citric acid study; some reward sizes that were tested did not qualify as a steady-state run (e.g. runs interrupted by a day with too great a fluctuation in wage rate or task performance, or a deviation from standard access schedule, or if the rat received a water supplement); some rats had time off from the study to recover from weight loss (e.g., after a too-small reward size was tested); some older rats exited the study early for age-related health reasons; and the naïve rats were added late in the study. The animals whose data are shown (N=7 across all analyses) were in each case all of the rats that had sufficient data for the respective analysis, by the indicated criteria. Details of how subjects in the cohort were screened for inclusion in each analysis, and the data from all animals whether or not included in any analysis, are provided in the data archive.

##### Measurement of reward size

Because the water volume of the reward was crucial to this study, we measured it in two ways. The left and right reward ports had separate water valves, which were chosen to be approximately matched for flow and then calibrated periodically to measure the (~linear) relationship between open time in ms and volume delivered in µl. The actual valve-open durations were recorded for each trial, and the expected reward (average ml/trial) was later computed on a per-day basis based on the number of rewards received on each side, the measured valve open times of those reward events, and the side-specific calibrated flow rates.

When rats lick a water tube to collect rewards, however, their tongues block the water flow part of the time, reducing the effective reward volume. To account for this, we also measured the total amount of water dispensed, by visually observing the change in water level of each station's individual water-supply syringe each day. Evaporative loss was prevented with loose-fitting covers. For most rats, the measured water consumption was about 75% of the volume predicted from calibrated flow, independent of the valve open time. This conversion factor was measured for each rat and used to correct calibrated measurements. We confirmed that the conclusions of the study were unaltered if the wage rate and intake were measured in 3-7 day blocks (as shown in Figure 1) instead of by day (Figure 3,4).

**Model Fitting Details**

The equation parameters were fit by an exhaustive grid search to minimize the residual squared error. This fit was deterministic. We started with a coarse grid uniformly sampling the plausible parameter value range ($\alpha_{24}$ from 8-80 ml/day; $\alpha_2$from 8-30 ml/day; $\beta$ 0.001-0.15). We evaluated the equations for every possible combination of parameter values, at each of the wage rates of the observed data points. We defined the residual error as the sum of the sum of squared differences between model and data for water intake H, divided by the degrees of freedom (N-1). We smoothed the error landscape by a narrow gaussian filter and then selected the parameter combination with the lowest residual error. The search grid was then re-centered on the selected parameter combination, the span of the search grid was contracted and the resolution increased by a factor of 2 (if 3 parameters were being fit) or 3 (if only two parameters, i.e. fitting data from only one schedule), excluding any illegal parameter values (bounds stated in equations). This was repeated at 4 progressively finer resolutions. For the four-parameter model we fit two parameters for each schedule in separate grid searches, and then used the schedule-specific parameters to predict held-out data from both schedules. Cross-validated residual errors of all models (including the fixed income model) were computed by leave-one-out cross-validation (predicting each data point by a model fit to all other data points).

**Task Variables Not Modeled**

The visual difficulty of the task was drawn randomly from a fixed distribution, such that some trials were harder than others. At the time of the rat's decision to initiate a trial, however, the *expected* difficulty is the same for every trial (the sum over coherence levels of the probability of that coherence times the subjective difficulty of that coherence). Both the expected reward and the uncertainty in reward are constant across trials and between reward size or schedule conditions. Our model seeks to predict the choice to initiate a trial, and therefore use a single parameter for disutility of work (β). The number of trials performed per session is in the range of hundreds to thousands, so the experienced average difficulty should be close to the theoretical expected difficulty in individual sessions. We did not experimentally vary the expected difficulty, and we changed the expected reward once a week or less often. Nevertheless, it is possible that the rat updates its estimates of α and β on the basis of random fluctuations in recently experienced reward and effort. This would be interesting to investigate in a larger dataset with sufficient power to constrain the additional parameters.

#### Statistics

This was a hypothesis-generating study. We do not generalize any conclusions to the population of rats as a whole, and we advisedly refrain from any statements of statistical significance. Rather we present the examples as existence proofs of the phenomena that motivated the model. The statistics we present are quantify the evidence for within-individual trends.

In Figure 1 each symbol represents the average empirically measured reward size and average trial number or water consumption over 4-7 consecutive days in a steady state condition. These blocks were identified by first finding contiguous runs of at least 4 days in a row with the same reward and schedule, excluding the first day after a change in reward or schedule, and then breaking up the run into nonoverlapping blocks of 4 days; if <4 days remained at the end of the run they were included in the last block. The purpose of this was to average out daily fluctuations (compare symbols in Figure 1 to the same data shown at one-day granularity in Figure 4). The reward conditions were interleaved on a counterbalanced schedule. We show data from all N=4 rats that were tested in at least 8 such blocks with reward sizes that spanned at least a 50µl range of values. Three other rats had too few time blocks or too narrow a range of reward size to support a meaningful analysis of correlation, but their statistics are also shown in the table for completeness (gray shading). Remaining rats in the study either had fewer than 3 independent time blocks or only one reward size (*w* range <3µl) and were not included in this analysis. These analysis details and exclusion criteria were decided after the data were known.

In panels A-D, our observation is that trial rate declined with increasing reward size, as opposed to being constant (as predicted by a fixed effort model) or increasing with reward size (as predicted if vigor increases with reward size). To quantify the evidence for this conclusion we used a one-tailed Spearman correlation vs. the null hypothesis that the correlation was $\rho \geq 0$.

In panels E-H, our observation is that water consumption increased with increasing reward size, as opposed to being constant (as predicted by the fixed income model, blue) or declining (as predicted by either fixed effort or vigor increasing with reward). To quantify the evidence for this conclusion, we used a one-tailed Spearman correlation vs. the null hypothesis that the correlation was $\rho \leq 0$. The fixed income model is the average observed consumption $k = \overline{H}$, blue lines in E-H. The number of trials predicted by this model is simply $L = k/w$, blue curves in A-D. We did not set a significance threshold because no binary significance decisions are planned. Any P values $<10^{-6}$ are reported as such because this exceeds the precision of the estimate of P from our data. Spearman correlation was used because the relationships are monotonic but not necessarily linear.

| | Time blocks | w range | k | w vs L | | | w vs H | | |
|---|---|---|---|---|---|---|---|---|---|
| Rat | N | µl/trial | ml/day | rho | $R^2$ | p | rho | $R^2$ | p |
| A | 12 | 68 | 17.1 | -0.94 | 0.88 | <1e-6 | 0.83 | 0.68 | 8.60e-04 |
| B | 10 | 54 | 18.4 | -0.96 | 0.93 | <1e-6 | 0.95 | 0.91 | <1e-6 |
| C | 11 | 84 | 17.3 | -0.98 | 0.96 | <1e-6 | 0.85 | 0.71 | 1.00e-03 |
| D | 8 | 81 | 20.0 | -0.98 | 0.95 | 2.00e-04 | 0.93 | 0.86 | 1.10e-03 |
| E | 9 | 16 | 15.2 | -0.68 | 0.47 | 2.50e-02 | 0.37 | 0.13 | 1.70e-01 |
| F | 5 | 12 | 15.9 | -0.8 | 0.64 | 6.70e-02 | 0.30 | 0.09 | 3.40e-01 |
| G | 3 | 37 | 19.7 | -1.00 | 1.00 | 1.70e-01 | 0.50 | 0.25 | 5.00e-01 |

**Table S1. Statistics supporting Figure 1, and data not shown (shaded)**

### Supplemental Results

#### Schedule dependence of water satiety

The rats from which the wage labor data were collected had occasionally received free water, but consumption was not measured. The value of this measurement became apparent only later when the model was developed. Therefore, we were unable to constrain the free parameter α in the model fits.

In a separate group of 6 female Long-Evans rats, however, we measured steady-state free water consumption on either a 24-hour or 2-hour access schedule (Supplemental Figure). Average free water consumption for these rats was 23.8±3.9 ml/day (mean±SD, N=6 animals) on 24-hour access, vs. 12.1±1.5 ml/day on 2-hour per day access. These data confirm that water consumption is schedule-dependent, and the values are consistent with the estimates of α estimated in our test rat population.

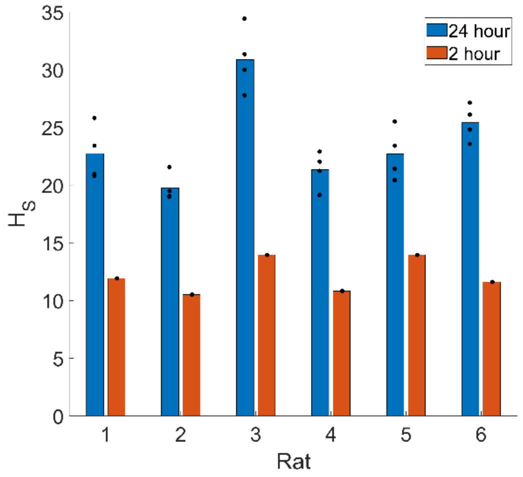


**SI Figure 1. Free water satiety depends on access schedule.** Water consumption was measured for singly housed rats over a span of at least four consecutive days with water access either 2 hours/day at the same time each day, or 24 hours/day. The 24 hour/day condition experiment was repeated four separate times, the 2 hour/day condition was tested once (black symbols).

#### Alternative utility equations

We presented the simplest equations sufficient to explain our data. In the future, however, richer datasets (such as those generated by the experiment suggested in Figure 6) may be sufficient to reveal additional dependencies. Therefore, alternative utility functions are noted here. Note that we have chosen notations for which marginal utility has a simple form, as this is the quantity directly relatable to time varying behavior or neural representations.

The first utility equation we tried for water had three parameters: $U_H(H) = \frac{\alpha}{1-\gamma} H^{1-\gamma} - \delta H$ , such that $MU_H(H) = \alpha H^{-\gamma} - \delta$. In this case, the parameter $\delta$ can be constrained by the satiety point, still leaving two free parameters. For a single-parameter equation one could also try a quadratic utility function such as $U_H(H) = H - \frac{\alpha}{2} H^2$, such that $MU_H(H) = 1 - \alpha H$. In this case the satiety point could be used to constrain the single parameter $\alpha$.

We initially assumed the aversiveness of labor would increase with the number of trials done (i.e. the $100^{th}$ pushup of the day is far more aversive than the first one). Thus, an alternative two-parameter equation for the utility of labor (c.f. Equation 3) could be $U_L(L) = \frac{\gamma}{2} L^2 - \beta L$, such that marginal disutility scales linearly with trial number: $MU_L(L) = \gamma L - \beta$; or even $U_L(L) = \frac{-1}{1+\gamma} L^{1+\gamma} - \beta L$, such that marginal disutility scales exponentially with trial number: $MU_L(L) = -L^\gamma - \beta$.

Finally, we have made the simplifying assumption that there are no cross-terms: the utility of water $U_H(H)$ does not depend on the amount of work $L$, and the disutility of labor $U_L(L)$ does not depend on the amount of water $H$. One could imagine circumstances in which these would interact, however, such as if the "work" were so physically strenuous as to dehydrate the animal.
